# A Multi-Omics Longitudinal Aging Dataset in Primary Human Fibroblasts with Mitochondrial Perturbations

**DOI:** 10.1101/2021.11.12.468448

**Authors:** Gabriel Sturm, Anna S Monzel, Kalpita R Karan, Jeremy Michelson, Sarah A. Ware, Andres Cardenas, Jue Lin, Céline Bris, Balaji Santhanam, Michael P Murphy, Morgan E Levine, Steve Horvath, Daniel W Belsky, Shuang Wang, Vincent Procaccio, Brett A. Kaufman, Michio Hirano, Martin Picard

## Abstract

Aging is a process of progressive change. In order to develop biological models of aging, longitudinal datasets with high temporal resolution are needed. Here we report a multi-omic longitudinal dataset for cultured primary human fibroblasts measured across their replicative lifespans. Fibroblasts were sourced from both healthy donors (n=6) and individuals with lifespan-shortening mitochondrial disease (n=3). The dataset includes cytological, bioenergetic, DNA methylation, gene expression, secreted proteins, mitochondrial DNA copy number and mutations, cell-free DNA, telomere length, and whole-genome sequencing data. This dataset enables the bridging of mechanistic processes of aging as outlined by the “hallmarks of aging”, with the descriptive characterization of aging such as epigenetic age clocks. Here we focus on bridging the gap for the hallmark mitochondrial metabolism. Our dataset includes measurement of healthy cells, and cells subjected to over a dozen experimental manipulations targeting oxidative phosphorylation (OxPhos), glycolysis, and glucocorticoid signaling, among others. These experiments provide opportunities to test how cellular energetics affect the biology of cellular aging. All data are publicly available at our webtool: https://columbia-picard.shinyapps.io/shinyapp-Lifespan_Study/

## Background & Summary

Aging is the major risk factor for all major diseases ^1^. In biological terms, aging involves progressive changes at multiple levels of molecular organization, including the genome ^2–4^, epigenome ^5–7^, transcriptome ^8^, proteome ^9^, secretome ^10^, and organs and organ-systems ^11^. Advances in aging biology have identified a set of molecular “hallmarks” or “pillars” thought to represent the root causes of aging-related declines in cellular- and organ-system integrity and subsequent disease, disability, and mortality ^12,13^. In parallel, recent expansion of omics technologies have enabled researchers to generate high-dimensional datasets across multiple modalities that illuminate the complex molecular landscape of biological aging. These data have been combined with machine-learning methods to develop molecular “clocks” that track chronological age and mortality risk with remarkable accuracy ^14,15^, and to model complex systems-level processes ^16,17^. The clocks and related measures make possible measurements of biological processes of aging in free-living humans. But their connections to the hallmarks of aging remain unclear. Research is needed to elucidate fundamental mechanisms that cause aging-related changes, and that drive aging-related declines in resilience and increased vulnerability to disease, disability, and mortality.

The ideal approach to meeting this research need would be to longitudinally monitor a population of individuals over their entire lifespan, taking regular measures of many metrics at frequent intervals. Such studies represent an important frontier in aging science ^18,19^. However, in addition to resource-constraints and participant-burden concerns that limit the frequency and depth of measurements, following a cohort of humans over their lifespan requires multiple generations of scientists and faces challenges of ever-changing techniques and technology for sample collection and analysis. One complementary strategy is to conduct lifespan studies of shorter-lived animals ^20,21^, although these too require substantial time, and may be limited in their translability to humans ^22^. Here we present a further complementary strategy: a “lifespan” study of cultured fibroblasts.

One contribution from cross-species work has been to delineate conserved genes, pathways, and hallmarks of aging across multiple experimental modalites ^12^. One major pathway surfacing as a critical and possibly primary driver of aging biology is mitochondrial metabolism ^23^. Omics-based discovery studies consistently converge on mitochondria as a major molecular signature of biological aging ^9,24^. Beyond providing ATP for all basic cellular functions, mitochondria produce signals that can trigger multiple hallmarks of aging ^25^. In humans ^26,27^ and animals ^28–30^, mitochondrial respiratory chain dysfunction dramatically shortens lifespan, further suggesting a primary causal role of mitochondria as a “timekeeper” and driver of the biological aging process.

To examine longitudinal aging trajectories and quantify the influence of mitochondria on canonical and exploratory aging markers in a human system, we generated a multi-omic, longitudinal dataset across the replicative lifespan of primary human fibroblasts from several healthy and disease donors (**Figure 1**). These data include genomic, epigenomic, transcriptomics, and protein-based measures along with bioenergetic and mitochondrial OxPhos measures. The relatively high temporal resolution of measurements allows for non-linear modeling of molecular recalibrations in primary human cells, as recently shown for DNA methylation in a pilot cellular lifespan study ^31^. In this cellular lifespan system, the rate of biological aging appears to proceed at a rate ∼40-140x faster than *in vivo* (i.e., in the human body), such that 200-300 days *in vitro* corresponds to multiple decades of human life ^31^. In addition to the rich descriptive data in multiple donors, this dataset includes experimental conditions with metabolic manipulations targeted to mitochondria, allowing investigators to directly test the influence of mitochondrial metabolism on human molecular aging signatures.

**Figure 1.**
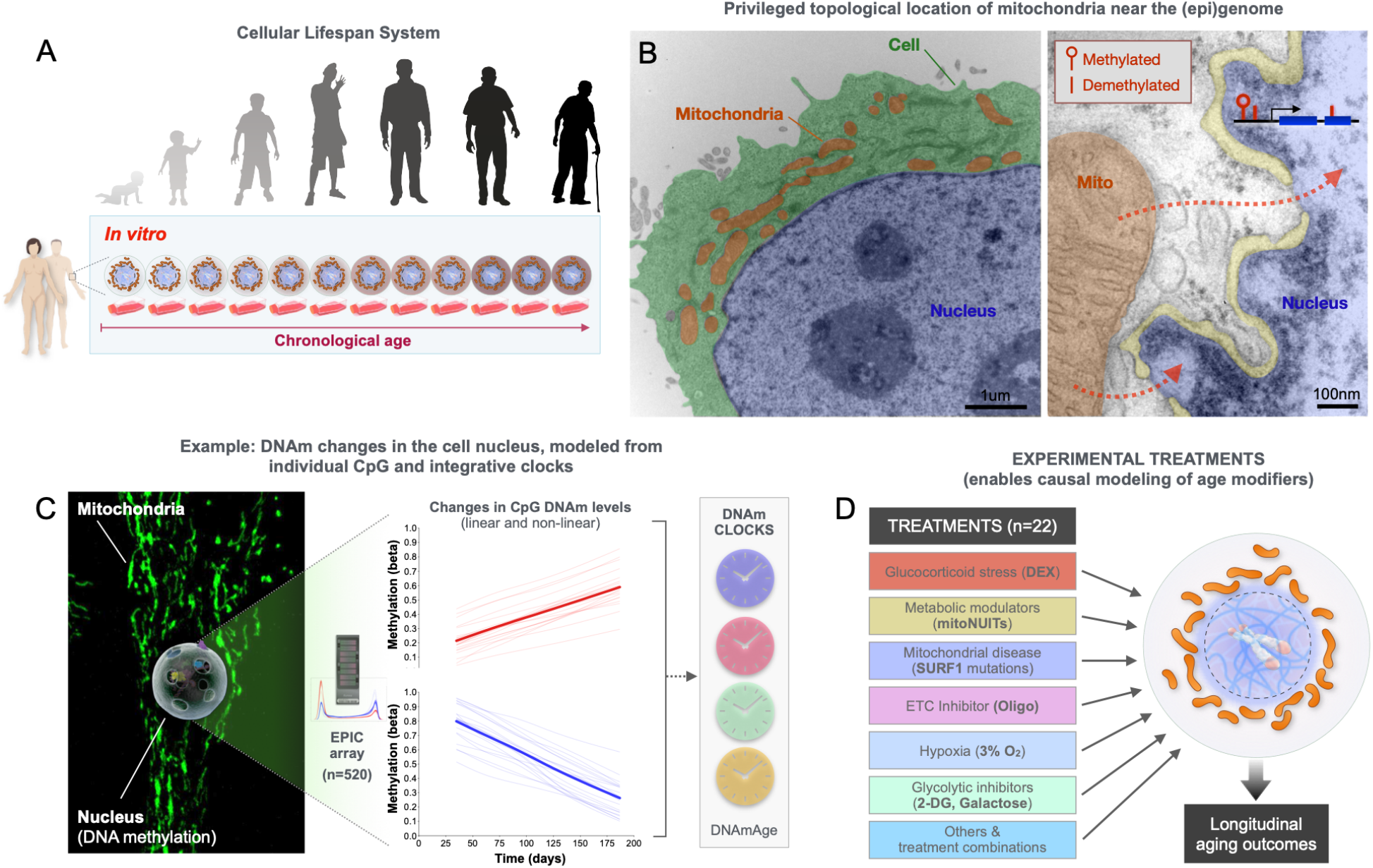
Biological and conceptual rationale for the *Cellular Lifespan Study*. (**A**) Primary human fibroblasts aged in culture (i.e., *in vitro*) recapitulate several, although not all, molecular hallmarks of human aging. Previous work with this replicative lifespan system showed that canonical age-related changes in DNA methylation (DNAm) in human tissues, such as hypermethylation of the *ELOVL2* gene promoter (cg16867657), global hypomethylation, and the rate of epigenetic aging captured by epigenetic clocks, are conserved, but occur at an accelerated rate in cultured primary human fibroblasts ^31^. This model provides a system to recapitulate and model some of the longitudinal changes in the cells of the same individuals, at high temporal resolution across the replicative lifespan. (**B**) Electron micrograph of a human cultured cell (*left*) and higher magnification view of the surface of the nucleus and nuclear envelope (yellow), with a neighboring mitochondrion. Arrows illustrate the diffusion path for soluble metabolic signals to reach the (epi)genome. Illustration modified from ^32^. (**C**) Micrograph of a whole fibroblast (HC2, P22, 103 days grown) with fluorescently-labeled mitochondria (MitoTracker green) surrounding the nucleus, from which DNA methylation can be measured using the EPIC array, quantified either at the single CpG level, or integrating data from multiple CpGs via different DNAm clocks. (**D**) Illustration of the experimental segment of the *Cellular Lifespan Study*, where specific signaling pathways (glucocorticoid signaling), OxPhos (oligomycin) and glycolytic pathways (no glucose, galactose, 2-deoxyglucose), respiratory chain defects (donors with SURF1 mutations), and other single and combinations of treatments were used to perturb selected metabolic pathways.

## Methods

### Tissue culture

Primary human dermal fibroblasts were obtained from commercial distributors or in our local clinic from 6 healthy control donors, and 3 donors with lethal SURF1 mutations (IRB #AAAB0483) that alter mitochondrial respiratory chain complex IV assembly and function ^33,34^, and lead to early death in affected patients ^35^ (see **Table 1** for descriptive information and distributor). SURF1-mutant fibroblasts were isolated from dermal punch biopsies of the forearm skin using standard procedures. After isolation, fibroblasts were cryopreserved in 10% DMSO (Sigma-Aldrich #D4540), 90% fetal bovine serum (FBS, Life Technologies #10437036) in liquid nitrogen. To avoid freeze-shock necrosis cells were frozen gradually in an isopropanol container (Thermofisher #5100-0001) at -80°C overnight before storage in liquid nitrogen. Cells were thawed at 37°C (<4min) and immediately transferred to 20ml of pre-warmed DMEM (Invitrogen #10567022).

**Table 1.** Cell line metadata. Demographic, tissue of origin, and genetic characteristics of the primar human fibroblast cell lines used in the *Cellular Lifespan Study* dataset. The dataset includes 6 healthy control donors, and 3 donors with lethal SURF1 mutations that alter mitochondrial respiratory chain complex IV assembly and function, and lead to early death in affected patients.

| Cell Line name | In-house name | Received From | Tissue Location | Genetic | SURF1 Mutation | Ethnicity | Sex | Donor Age (years) | Passage | Cat # |
| --- | --- | --- | --- | --- | --- | --- | --- | --- | --- | --- |
| HC1 | hFB12 | Lifeline Cell Technology | Dermal Breast | Normal | N/A | Caucasian | male | 18 | 1 | FC-0024 Lot # 03099 |
| HC2 | hFB13 | LifeLine Cell Technology | Dermal Breast | Normal | N/A | Caucasian | female | 18 | 1 | FC-0024 Lot # 00967 |
| HC3 | hFB14 | Corriel Institute | Foreskin | Normal | N/A | Black | male | 0 | 4 | AG01439 |
| HC4 | hFB11 | Hirano Lab | Dermal Upper-Arm Skin | Normal | N/A | N/A | male | 3 | 4 |  |
| HC5 | hFB1 | Hirano Lab | Dermal Upper-Arm Skin | Normal | N/A | Caucasian | male | 29 | 3 |  |
| HC6 | hFB2 | Hirano Lab | Dermal Upper-Arm Skin | Normal | N/A | Caucasian | female | 26 | 4 |  |
| SURF1_1 | hFB6 | Hirano Lab | Dermal Upper-Arm Skin | SURF1 Mutation | c.518_519del (p.S173Cfs*7)<br>c.845_846del (p.S282Cfs*7) | N/A | male | 0.25 | 7 |  |
| SURF1_2 | hFB7 | Hirano Lab | Dermal Upper-Arm Skin | SURF1 Mutation | c.247_248insCTGC (p.R83Pfs*7)<br>c.313_321del (p.L105_A107del) | N/A | male | 11 | 5 |  |
| SURF1_3 | hFB8 | Hirano Lab | Dermal Upper-Arm Skin | SURF1 Mutation | Homozygous c.313_321del (p.L105_A107del) | N/A | female | 9 | 9 |  |

For replicative lifespan studies, cells were cultured in T175 flasks (Eppendorf #0030712129) at standard 5% CO_2_ and atmospheric O_2_ at 37°C in DMEM (5.5 mM glucose) supplemented with 10% FBS (Thermofisher #10437010), 50 μg/ml uridine (Sigma-Aldrich #U6381), 1% MEM non-essential amino acids (Life Technologies #11140050), 10 μM palmitate (Sigma-Aldrich #P9767) conjugated to 1.7 μM BSA (Sigma-Aldrich #A8806), and 0.001% DMSO (treatment-matched, Sigma-Aldrich #D4540). Cells were passaged approximately every 6 days (+/- 1 day), with decreasing passaging frequency as cells enter quiescence, for up to 270 days.

Brightfield microscopy images at 10x and 20x magnification were taken before each passaged using inverted phase-contrast microscope (Fisher Scientific #11350119). All images except those of Phase IV can be downloaded at: https://figshare.com/articles/dataset/Brightfield_Images_for_Cellular_Lifespan_Study/18444731.

Cell counts, volume and proportion of cell death were determined in duplicates (CV <10%) and averaged at each passage using the Countess II Automated Cell Counter (Thermofisher #A27977). To determine the number of cells to plate at each passage, growth rates from the previous passage were used, pre-calculating the expected number cells needed to reach ∼90% confluency (∼2.5 million cells) by the next passage, ensuring relatively similar confluence at the time of harvesting for molecular analyses. Cells were never plated below 200,000 cells or above million cells to avoid plating artifacts of isolation or contact inhibition, respectively. However, some differences in cell density between early and late passages were unavoidable. Study measurements and treatment began after 15-days of culturing post-thaw to allow for adjustment to the *in vitro* environment. Individual cell lines from each donor were grown until they exhibited less than one population doubling over a 30-day period, at which point the cell line was terminated, reflecting the end of the lifespan. The hayflick limit was calculated as the total number of population doublings reached by the end of each experiment.

### Mycoplasma testing

Mycoplasma testing was performed according to the manufacturer’s instructions (R&D Systems #CUL001B) on 100 media samples at the end of lifespan for each treatment and cell line used. All tests were negative.

### Bioenergetic parameters and calculations of metabolic rate

Bioenergetic parameters were measured using the XFe96 Seahorse extracellular flux analyzer (Agilent) ^36,37^. Oxygen consumption rate (OCR) and extracellular acidification rate (ECAR, i.e. pH change, was measured over a confluent cell monolayer. Cells were plated for Seahorse measurement every 3 passages (∼15 days) with 10-12 wells plated per a treatment group. Each well of the Seahorse 96-well plate was plated with 20,000 cells and incubated overnight under standard growth conditions, following the manufacturer’s instructions, including a plate wash with complete Seahorse XF Assay media. The complete XF media contains no pH buffers and was supplemented with 5.5 mM glucose, 1 mM pyruvate, 1 mM glutamine, 50 μg/ml uridine, 10 μM palmitate conjugated to 1.7 μM BSA, to ensure that cells have access to a variety of energetic substrates. After a wash with XF media, the plate was incubated with XF media in a non-CO_2_ incubator for one hour to equilibrate temperature and atmospheric gases before the assay.

Different respiratory states were assessed using the MitoStress Test ^38^. Basal respiration, ATP turnover, proton leak, coupling efficiency, maximum respiration rate, respiratory control ratio, spare respiratory capacity, and non-mitochondrial respiration were all determined by the sequential additions of the ATP synthase inhibitor oligomycin (final concentration: 1 μM), the protonophore uncoupler FCCP (4 μM), and the electron transport chain Complex I and III inhibitors, rotenone and antimycin A (1 μM). The optimal concentration for the uncoupler FCCP yielding maximal uncoupled respiration was determined based on a titration performed on young healthy fibroblasts (data not shown). The final injection included Hoechst nuclear fluorescent stain (Thermofisher #62249) to allow for automatic cell counting. After each run, cell nuclei density in each well were counted using the Cytation1 Cell Imager (BioTek) and raw bioenergetic measurements were normalized to relative cell counts on a per-well basis. This normalization method was selected due to reduced well-to-well variability by approximately half compared to other normalization techniques (e.g., normalization to ug of protein).

ATP production rates from oxidative phosphorylation (OxPhos, *J*_ATP-OxPhos_) and glycolysis (*J*_ATP-Glyc_), as well as total cellular ATP production and consumption (*J*_ATP-Total_) were estimated using the method described by Mookerjee et al. ^36^. Briefly, the method relies on the phosphate-to-oxygen (P/O) ratios of OxPhos and glycolysis, using oxygen consumption and proton production rates (PPR) as input variables. The same constants were used for all estimations, assuming glucose as the predominant carbon source, and constant coupling efficiency. Changes in substrate consumption along the lifespan would require parallel assessments of metabolic flux to resolve, and assuming the same major substrate across the lifespan and treatment conditions could have a minor influence on calculated ATP production rates that are not reflected in the ATP-related metrics in this bioenergetics data. All raw seahorse data files and analysis scripts are available at https://github.com/gav-sturm/Cellular_Lifespan_Study/tree/main/Seahorse.

### mtDNA next-generation sequencing and eKLIPse analysis

The entire mtDNA was amplified in two overlapping fragments using a combination of mtDNA primers. The primer pairs used for PCR amplicons were tested first on Rho zero cells devoid of mtDNA to remove nuclear-encoded mitochondrial pseudogene (NUMTS) amplification

◦ PCR1: 5’-AACCAAACCCCAAAGACACC-3’ and 5’- GCCAATAATGACGTGAAGTCC-3’
◦ PCR2: 5’-TCCCACTCCTAAACACATCC-3’ and 5’- TTTATGGGGTGATGTGAGCC-3’

Long-range PCR was performed with the Kapa Long Range DNA polymerase according to the manufacturer’s recommendations (Kapa Biosystems, Boston, MA, USA), with 0.5µM of each primer and 20ng of DNA. The PCR products were analyzed on a 1% agarose gel electrophoresis.

NGS Libraries were generated using an enzymatic DNA fragmentation approach using Ion Xpress Plus Fragment Library Kit. Library were diluted at 100 pM before sequencing and pooled by a maximum of 25 samples.

Sequencing was performed using an Ion Torrent S5XL platform using Ion 540 chip™. Signal processing and base calling were done by the pre-processing embedded pipeline. Demultiplexed reads were mapped according to the mtDNA reference sequence (NC_012920.1) before being analysed with a dedicated homemade pipeline including eKLIPse ^39^ (https://github.com/dooguypapua/eKLIPse) using the following settings:

◦ Read threshold: min Quality=20 | min length =100bp
◦ Soft-Clipping threshold: Read threshold: Min soft-clipped length =25pb | Min mapped Part=20 bp
◦ BLAST thresholds: min=1 | id=80 | cov=70 | gapopen=0 | gapext=2
◦ Downsampling: No

### mtDNA copy number

Cellular mtDNA content was quantified by qPCR on the same genomic material used for other DNA-based measurements. Duplex qPCR reactions with Taqman chemistry were used to simultaneously quantify mitochondrial (mtDNA, ND1) and nuclear (nDNA, B2M) amplicons, as described previously ^40^. The reaction mixture included TaqMan Universal Master mix fast (life technologies #4444964), 300nM of custom design primers and 100nM probes:

◦ ND1-Fwd: 5’-GAGCGATGGTGAGAGCTAAGGT-3’
◦ ND1-Rev: 5’-CCCTAAAACCCGCCACATCT-3’
◦ ND1-Probe: 5’-HEX-CCATCACCCTCTACATCACCGCCC-3IABkFQ-3’
◦ B2M-Fwd: 5’-CCAGCAGAGAATGGAAAGTCAA-3’
◦ B2M-Rev: 5’-TCTCTCTCCATTCTTCAGTAAGTCAACT-3’
◦ B2M-Probe: 5’-FAM-ATGTGTCTGGGTTTCATCCATCCGACA-3IABkFQ-3’

The samples were cycled in a QuantStudio 7 flex qPCR instrument (Applied Biosystems) at 50°C for 2 min, 95°C for 20 sec, 95°C for 1min, 60°C for 20 sec, for 40 cycles. qPCR reactions were setup in triplicates in 384 well qPCR plates using a liquid handling station (epMotion5073, Eppendorf), in volumes of 20ul (12ul mastermix, 8ul template). Triplicate values for each sample were averaged for mtDNA and nDNA. Ct values >33 were discarded. For triplicates with a coefficient of variation (C.V.) > 0.02, the triplicates were individually examined and outlier values removed where appropriate (e.g., >2 standard deviations above the mean), with the remaining duplicates were used. The final cutoff for acceptable values was set at a C.V.= 0.1 (10%); samples with a C.V. > 0.1 were discarded. A standard curve along with positive and negative controls were included on each of the seven plates to assess plate-to-plate variability and ensure that values fell within measurement range. The final mtDNAcn was derived using the ΔCt method, calculated by subtracting the average mtDNA Ct from the average nDNA Ct. mtDNAcn was calculated as 2^ΔCt^ x 2 (to account for the diploid nature of the reference nuclear genome), yielding the estimated number of mtDNA copies per cell.

### RNA sequencing

Total genomic RNA was isolated for 360 samples every ∼11 days across the cellular lifespan for control lines and selected treatments. RNA was stabilized using TRIzol (Invitrogen #15596026) and stored at -80°C until extraction as a single batch. RNA was extracted on-column using a RNeasy kit (Qiagen #74104), DNase treated according to the manufacturer’s instructions, and quantified using the QUBIT high sensitivity kit (Thermofisher #Q32852). RNA integrity was quantified on Bioanalyzer (Agilent RNA nano kit 6000, #5067-1511) and Nanodrop 2000. Of the 360 samples, 352 had an RNA integrity number (RIN) score >8.0, a A260/A280 ratio between 1.8-2.2, and no detectable levels of DNA. For cDNA library preparation, 1,500ng of RNA at 50ng/μl was processed using Ribo-Zero Gold purification (QIAseq FastSelect -rRNA HMR Kit #334387) and NEBNext^®^ Ultra^™^ II RNA Library Prep Kit (Illumina #E7770L). cDNA was sequenced using paired-end 150bp chemistry on a HiSeq 4000 instrument (Illumina, single index, 10 samples/lane, Genewiz Inc). Sequencing depth was on average 40 million reads per sample. Post-sequencing QC (multiQC, v1.8) excluded 6 more samples, for a final sample set of 345. Sequenced reads were then aligned using the pseudoalignment tool *kallisto* v0.44.0 ^41^. This data was imported using txi import (‘tximport’, v1.18.0, length-scaled TPM), and vst normalized (‘DEseq2’, v1.30.1).

### DNA methylation

Global DNA methylation was measured on 512 samples using the Illumina EPIC microarray (Illumina, San Diego). Arrays were run at the UCLA Neuroscience Genomic Core (UNGC). DNA was extracted using the DNeasy kit (Qiagen #69506) according to the manufacturer’s protocol and quantified using QUBIT broad range kit (Thermofisher #Q32852). At least 375 ng of DNA was submitted in 30 µl of ddH_2_O to UNGC for bisulfite conversion and hybridization using the Infinium Methylation EPIC BeadChip kit. Samples with DNA below 12.5 ng/ul (∼90 of 512 samples) were concentrated using SpeedVac Vacuum Concentrator (Thermofisher #SPD1030A-115) for <1 hour. Sample positions were randomized across six assay plates to avoid systematic batch variation effects on group or time-based comparisons.

All DNA methylation data was processed in R (v4.0.2), using the ‘minfi’ package (v1.36.0). Quality control preprocessing was applied by checking for correct sex prediction, probe quality, sample intensities, and excluding SNPs and non-CpG probes. Of the 512 samples, 22 failed quality control and were excluded from further analysis, yielding a final analytical sample of n=479. Data was then normalized using functional normalization (Fun Norm). Using the R package ‘sva’ (v3.12.0), both RCP and ComBat adjustments were applied to correct for probe-type and plate bias, respectively. After quality control, DNAm levels were quantified as beta values for 865,817 CpG sites.

### DNA methylation clocks and related measures

We used DNA methylation data to calculate a series of measures broadly known as epigenetic clocks ^15^. We computed four clocks designed to predict the chronological age of the donor, Horvath1 (i.e. PanTissue clock) ^5^, Horvath2 (i.e. Skin&Blood clock) ^42^, Hannum ^43^, and PedBE ^44^ clocks; two clocks designed to predict mortality, the PhenoAge ^45^ and GrimAge ^46^ clocks; a clock to measure telomere length, DNAmTL ^47^; a clock designed to measure mitotic age, MiAge ^48^; a clock trained to predict cellular senescence, DNAmSen ^49^, and two DNA methylation measure of the rate of deterioration in physiological integrity, DunedinPoAm ^17^, and DundedinPACE ^50^.

For the Horvath, Hannum, PhenoAge, GrimAge, and DNAmTL clocks, this dataset includes both the original versions of these clocks, calculated using the online calculator hosted by the Horvath Lab (https://dnamage.genetics.ucla.edu/new) and versions developed using the methods proposed in Higgins-Chen et al. (https://github.com/MorganLevineLab/PC-Clocks) ^51^. Briefly, this method replaces the clock’s individual illumina probe measurements (5-500 CpGs) with the shared extracted variances among genome-wide CpGs from principal components (PC), yielding the PC-adjusted DNAmAges for each clock. Chronological age values used in the calculations of these clocks were the ages of the donors at the time of sampling. The MiAge clock was computed using the software published by in ^48^ (http://www.columbia.edu/~sw2206/softwares.htm). The Pace of Aging clocks, DunedinPoAm and DunedinPACE, were computed using the software published in ^17,50^ (https://github.com/danbelsky/DunedinPACE).

### Whole genome sequencing (WGS)

Total genomic DNA was isolated for 94 samples across cellular lifespan using column based DNeasy blood and tissue kit (Qiagen #69504) and quantified using Qubit dsBR assay (ThermoFisher #Q32850). Sample quality assessment, library preparation, whole genome sequencing and data pre-processing was performed by Genewiz using standard Illumina workflow. Briefly, WGS paired-end (PE) reads with 2×150bp configuration were obtained from Illumina HiSeq platform and processed using SAMtools (v1.2) and BaseSpace workflow (v7.0). PE reads were aligned to hg19 genome reference (UCSC) using Isaac aligner (v04.17.06.15) and BAM files were generated. Duplicate reads were identified and filtered using Picard tools (GATK). More than 80% of the bases were of high quality with a score >Q30. Mean depth of sequencing coverage was >20x with more than 90% of the genome covered at least 10 times. Variant calling from the entire genome was performed using Strelka germline variant caller (v2.8) for small variants including single nucleotide variants (SNVs) and insertion/deletion (Indels) and structural variants (SVs) were identified using Manta (v1.1.1). WGS data is available from the authors upon request.

### Telomere Length

Relative telomere length was evaluated on the same genomic material used for other DNA-based measurements. Measurements were performed by qPCR and expressed as the ratio of telomere to single-copy gene abundance (T/S ratio), as previously described ^52,53^. The reaction mixture included 20 mM Tris-HCl, pH 8.4; 50 mM KCl; 200 µM each dNTP; 1% DMSO; 0.4x Syber Green I; 22 ng E. coli DNA; 0.4 units of Platinum Taq DNA polymerase (Thermo Fisher Scientific #10966018) with custom design primers. The primers utilized were the following: i) For the telomere (T) PCR: tel1b [5’-CGGTTT(GTTTGG)5GTT-3’], used at a final concentration of 100 nM, and tel2b [5’-GGCTTG(CCTTAC)5CCT-3’], used at a final concentration of 900 nM; ii) For the single-copy gene (S, human beta-globin) PCR: HBG1 Fwd:[5’ GCTTCTGACACAACTGTGTTCACTAGC-3’], used at a final concentration of 300 nM, and HBG2 Rev: [5’-CACCAACTTCATCCACGTTCACC-3’], used at a final concentration of 700 nM. Approximately 6.6 ng of DNA template was added per 11 μL of the reaction mixture. A standard curve of human genomic DNA from buffy coat (Sigma-Aldrich #11691112001) along with positive and negative controls were included on each of plates to assess plate-to-plate variability and ensure that values fell within measurement range. The qPCR reactions were performed in triplicate using a LightCycler 480 qPCR instrument (Roche) using the following thermocycling conditions: i) For the telomere PCR: 96°C for 1 min, one cycle; 96°C for 1 sec, 54°C for 60 sec with fluorescence data collection, 30 cycles; ii) For the single-copy gene PCR: 96°C for 1 min, one cycle; 95°C for 15 sec, 58°C for 1 sec, 72°C for 20 sec, 8 cycles; 96°C for 1 sec, 58°C for 1 sec, 72°C for 20 sec, 83°C for 5 sec with data collection, 35 cycles. Triplicate values for each sample were averaged of T and S were used to calculate the T/S ratios after a Dixon’s Q test for outlier removal. T/S ratio for each sample was measured twice. For duplicates with C.V. > 0.07 (7%), the sample was run a third time and the two closest values were used. Only 5% of the samples (25 of 512 samples) had a C.V. > 0.01 after the third measurement, and the inter-assay C.V. = 0.03 ± 0.043. Telomere length assay for the entire study were performed using the same lots of reagents. Lab personnel who performed the assays were provided with de-identified samples and were blind to other data.

### Cytokines

Two multiplex fluorescence-based arrays were custom-designed with selected cytokines and chemokines based on human age-related proteins. Analytes were selected based on reported correlations of their levels in human plasma with chronological age ^54^, and based on their availability on R&D custom Luminex arrays (R&D, Luminex Human Discovery Assay (33-Plex) LXSAHM-33 and LXSAHM-15, http://biotechne.com/l/rl/YyZYM7n3). Cell media samples were collected at selected passages across cellular lifespan and frozen at -80°C until analysed as a single batch. Thawed samples were centrifuged at 500g for 5 min and supernatant transferred to a new tube. Media samples were ran undiluted, and the plates were incubated, washed, and signal intensity quantified on a Luminex 200 instrument (Luminex, USA) as per the manufacturer’s instructions. Positive (>200 days aged healthy fibroblast) and negative controls (fresh untreated media) samples were used in duplicates on each plate to quantify and adjust for batch variations. Data was fitted and final values interpolated from a standard curve in xPONENT v4.2. Cytokine concentrations were then normalized to the number of cells counted at the time of collection to produce estimates of cytokine production on a per-cell basis. Quantification of two selected cytokines, interleukin 6 (IL-6, Abcam#ab229434) and growth differentiation factor 15 (GDF15, R&D#DGD150), were repeated using enzyme-linked immunosorbent assays (ELISAs), according to the manufacturer’s instructions.

### Automated cell-free DNA isolation from cell culture media

Total cell-free DNA (cf-DNA) was isolated from cell culture media using a previously published automated, high throughput methodology ^55^. In brief, thawed cell culture media was centrifued at 1000g for 5min. 75 µL of supernatant was then dispensed into a clean 96 deep-well plate (Thermo Fisher, cat#95040450) using a Freedom EVO 150 automated liquid handler (Tecan, cat#10641150). 5.7 µL of 20 mg/mL Proteinase K (VWR, cat#97062-242) and 7.5 µL of 20% sodium dodecyl sulfate (Boston BioProducts, cat#BM-230) were subsequently dispensed into each well using a HandyStep digital repeater pipette (BrandTech Scientific, cat#705002). The plate was sealed with an adhesive PCR seal (Thermo Fisher Scientific, cat#AB0558) and covered with generic packaging tape. The plate was centrifuged at 500 x *g* for 1 minute (min), then incubated in an New Brunswick Innova 44 incubator shaker (Eppendorf, cat#M1282-0000) at 70°C for 16 hours. Following the overnight incubation, the plate was left at room temperature for 15 min and centrifuged again. After the seal was removed, 125 µL of MagMAX Cell Free DNA Lysis/Binding Solution (Thermo Fisher cat#AM8500) and 5 µL of Dynabeads MyOne Silane magnetic beads (Thermo Fisher cat#37002D) were dispensed into each well using the repeater pipette. The plate was loaded onto a KingFisher Presto (Thermo Fisher, cat#5400830) magnetic particle processor to begin the extraction process. The DNA-bound magnetic beads were washed three times with the following solutions: 1) 265 µL of MagMAX Cell Free DNA Wash Solution (Thermo Fisher, cat#A33601), 2) 475 µL of 80% ethanol, and 3) 200 µL of 80% ethanol. The cf-DNA was resuspended in 60 µL of MagMAX Cell Free DNA Elution Solution (Thermo Fisher, cat#33602) and stored at -20°C in a sealed 96-well PCR plate (Genesee Scientific, cat#24-302).

### Quantification of cf-mtDNA and cf-nDNA abundance

*qPCR*: cf-mtDNA and cf-nDNA levels were measured simultaneously by qPCR. Taqman-based duplex qPCR reactions targeted mitochondrial-encoded ND1 and nuclear-encoded B2M sequences as described previously ^55–57^. Each gene assay contained two primers and a fluorescent probe and were assembled as a 20X working solution according to the manufacturer’s recommendations (Integrated DNA Technologies). The assay sequences are:

◦ ND1-Fwd: 5’-GAGCGATGGTGAGAGCTAAGGT-3’
◦ ND1-Rev: 5’-CCCTAAAACCCGCCACATCT-3’
◦ ND1-Probe: 5’-5HEX/CCATCACCC/ZEN/TCTACATCACCGCCC/2IABkGQ-3’
◦ B2M-Fwd: 5’-TCTCTCTCCATTCTTCAGTAAGTCAACT-3’
◦ B2M-Rev: 5’-CCAGCAGAGAATGGAAAGTCAA-3’
◦ B2M-Probe: 5’-56FAM-ATGTGTCTG-ZEN-GGTTTCATCCATCCGACCA-3IABkFQ-3’

Each reaction contained 4 µL of 2X Luna Universal qPCR Master Mix (New England Biolabs, cat#M3003E), 0.4 µL of each 20X primer assay, and 3.2 µL of template cf-DNA for a final volume of 8 µL. The qPCR reactions were performed in triplicate using a QuantStudio 5 Real-time PCR System (Thermo Fisher, cat#A34322) using the following thermocycling conditions: 95°C for 20 s followed by 40 cycles of 95°C for 1 s, 63°C for 20 s, and 60°C for 20 s. Serial dilutions of pooled human placenta DNA were used as a standard curve.

### Digital PCR (dPCR)

mtDNA and nDNA copy number (copies/µL) of the standard curve used in cf-mtDNA/cf-nDNA assessment were measured separately using singleplex ND1 and B2M assays using a QuantStudio 3D Digital PCR System and associated reagents (Thermo Fisher, cat#A29154) according to the manufacturer’s protocol. The values obtained for the standard curve were used to calculate the copy number for the experimental samples. All reactions were performed in duplicate (two chips). Because the same standard curve was used on all plates, its copy number was applied uniformly to all qPCR plates.

## Data Records

This multi-omics cellular lifespan dataset includes longitudinal data across 13 major biological outcomes, including cytological measures (growth rate, cell size), cellular bioenergetics (respiratory capacity, total energy consumption), transcriptomics (bulk RNA-seq), DNA methylation (EPIC array), whole genome sequencing (WGS), secreted factors (cytokines, cell-free DNA), and others (**Figure 2-3** and **Table 3**). These outcomes differ by the number of samples, repeat experiments, and length of timecourses for each donor line and treatment group (**Supplementary File 2** and **Table 4**). All expected hallmarks of cellular aging were observed in this model, including the upregulation of quiescence and senescence markers, and downregulation of genes associated with cell division (**Figure 4**).

**Figure 2.**
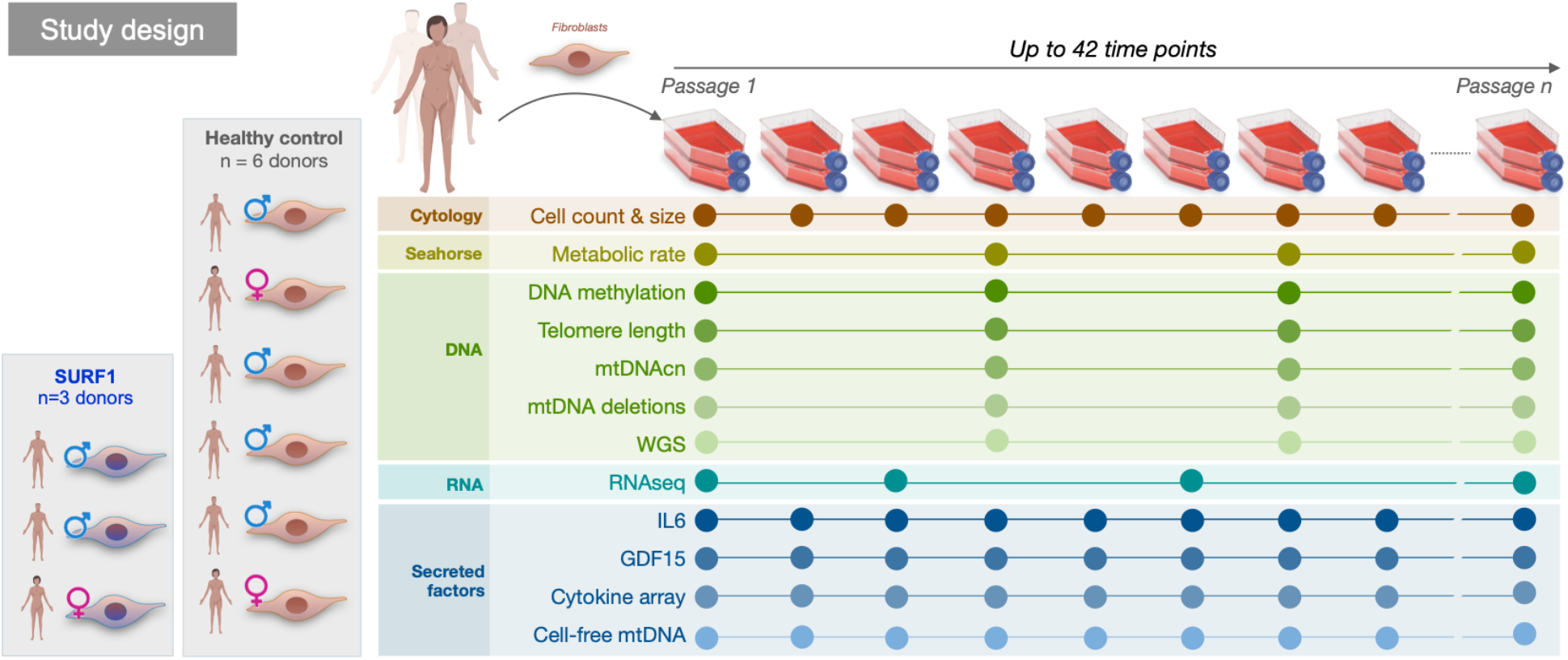
Overview of the study design and temporality of outcome measures. The outcome measures listed on the left were collected longitudinally at multiple time points following different periodicity, determined by resource or cell number constraints. The duration and periodicity of measurements vary by experimental conditions and cell line. See Tables 1 and 2 for details.

**Figure 3.**
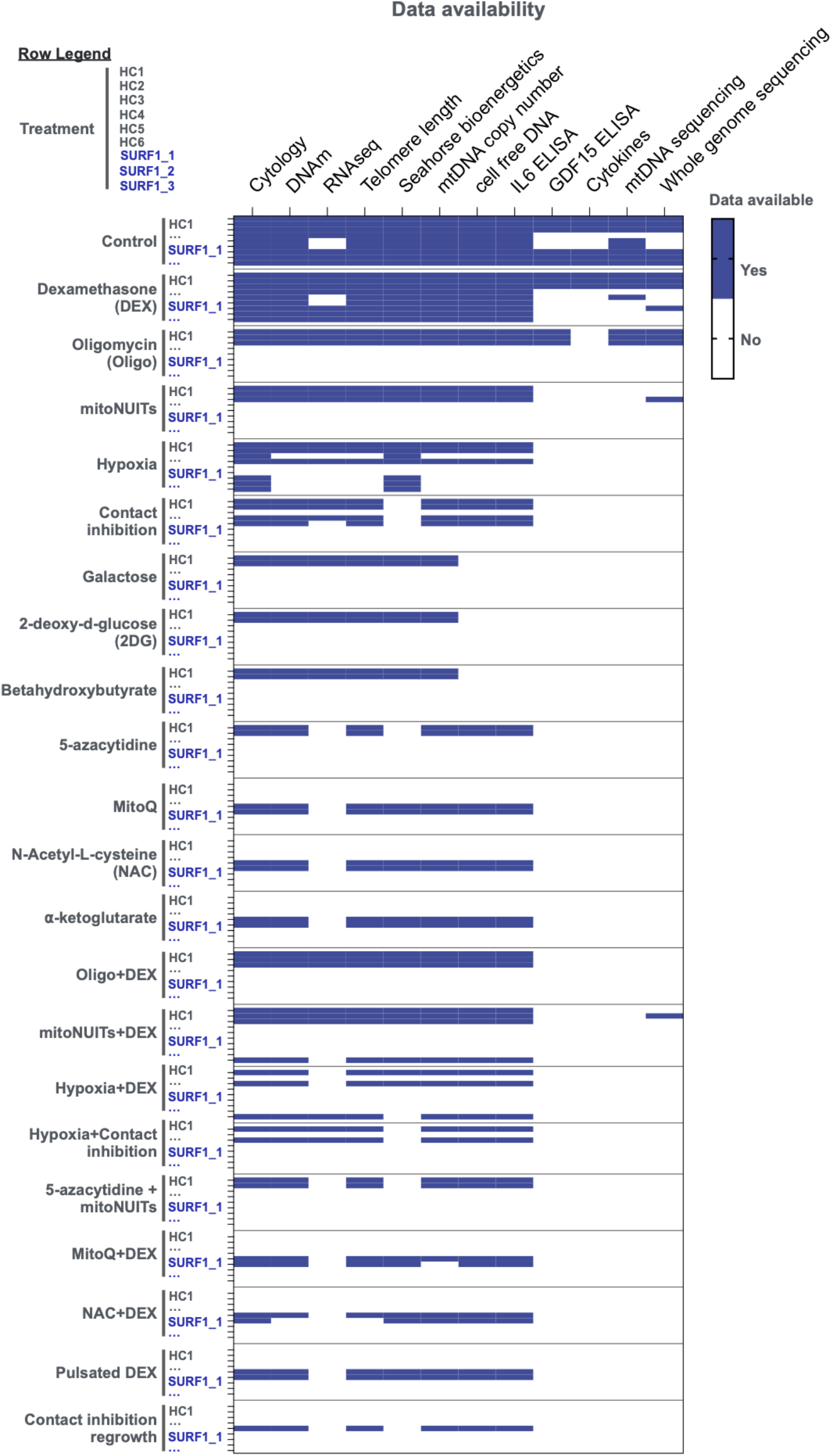
Data availability for each measured parameter. Heatmap of sample availability for each parameter (columns) associated with a given donor cell line and treatment (rows).

**Figure 4.**
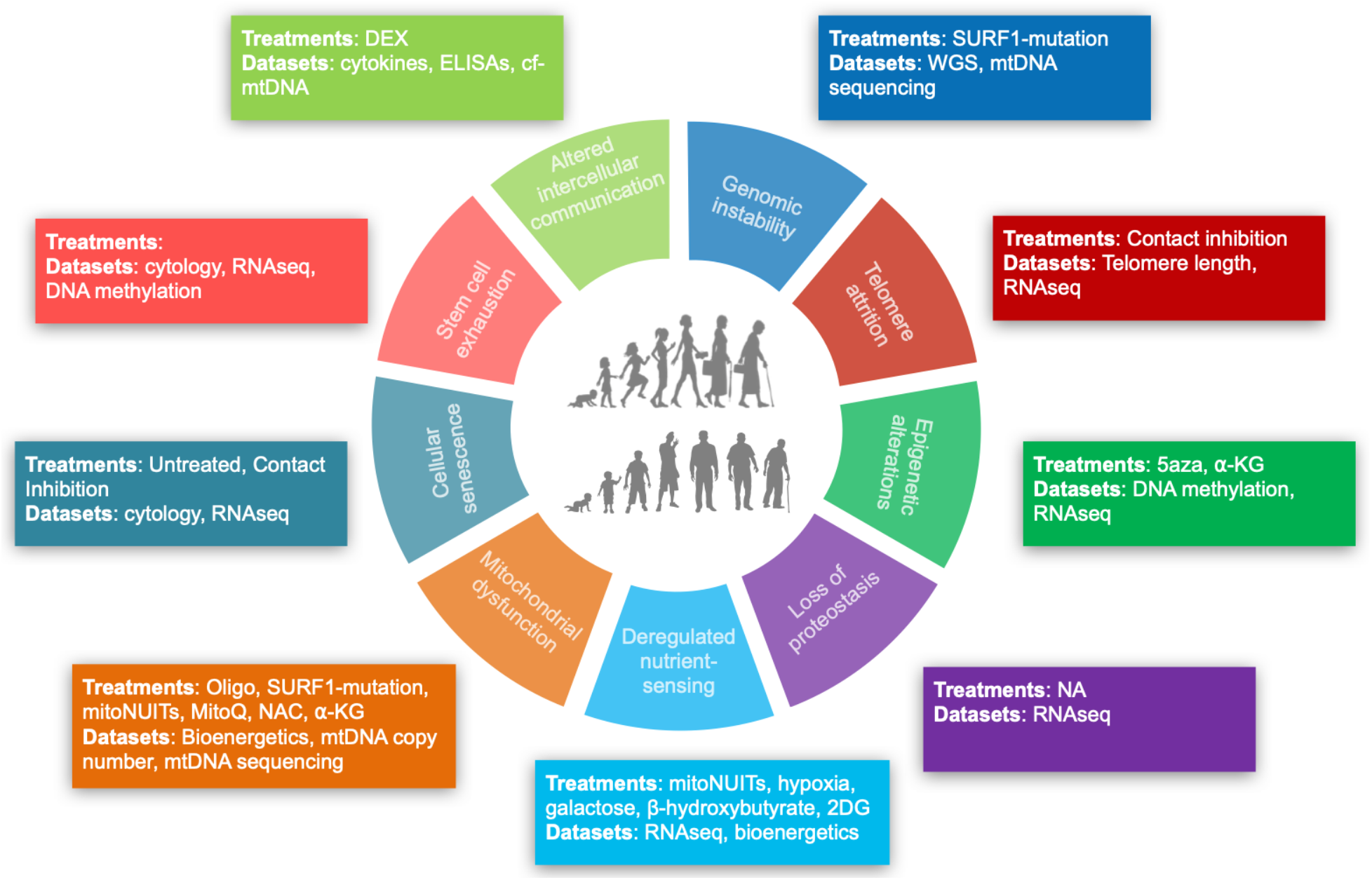
Intersection between the *Cellular Lifespan Study* conditions and the hallmarks of aging. Schematic of the hallmarks of aging adapted from López-Otín et al. ^12^, annotated with the treatments and datasets that directly or indirectly enable to longitudinally investigate their interplay with molecular features of cellular aging in aging cultured primary human fibroblasts.

**Table 1** contains information about all cell lines used, including the biopsy site, sex and age of the donor, as well as known clinical and genetic information.

**Table 2** indicates the cell lines with available data for each of the 22 experimental treatments, which are described below.

**Table 2.**
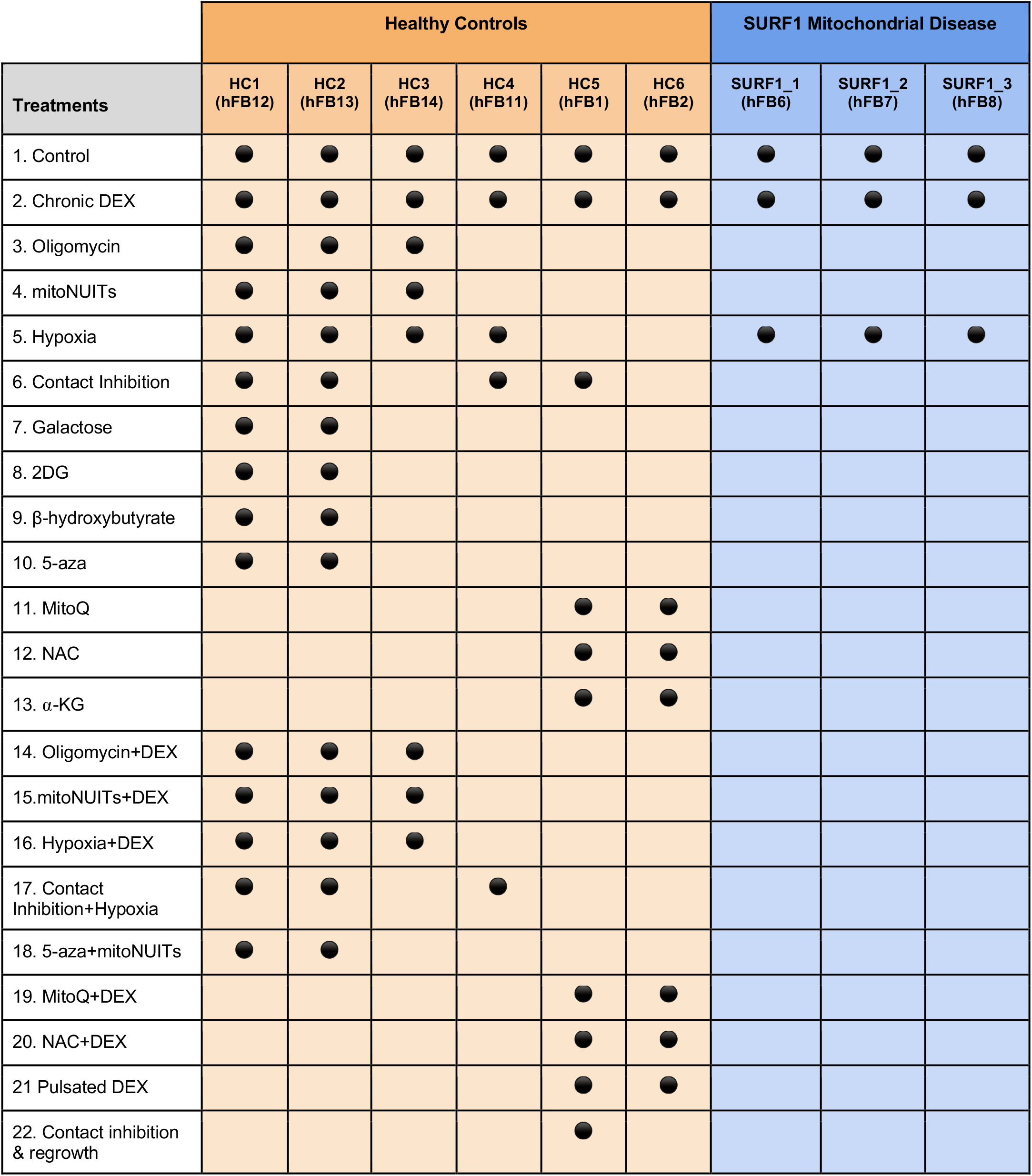
Cell lines and treatments for the *Cellular Lifespan Study*. Experimental conditions, or absence thereof, applied to each primary human fibroblast cell line.

**Table 3** lists the assessed biological outcomes along with their dimensionality (number of parameters quantified), the number of samples available across all cell lines, their temporal frequency, and the total size or number of datapoints. For example, there is available transcriptomic (i.e., gene expression) data quantified by RNA sequencing, which includes read counts for each annotated gene expressed as transcripts per million (TPM), on 360 samples collected on average every 11 days (min: 5 days, max 15 days), and each cell line has on average 8 timepoints (min:2, max:19). The transcriptomic dataset includes a total of 12.6B datapoints. Although multiple treatments were used to perturb bioenergetic and endocrine pathways, the dataset is most extensive and is of highest temporal resolution for cell lines from healthy controls.

**Table 3.**
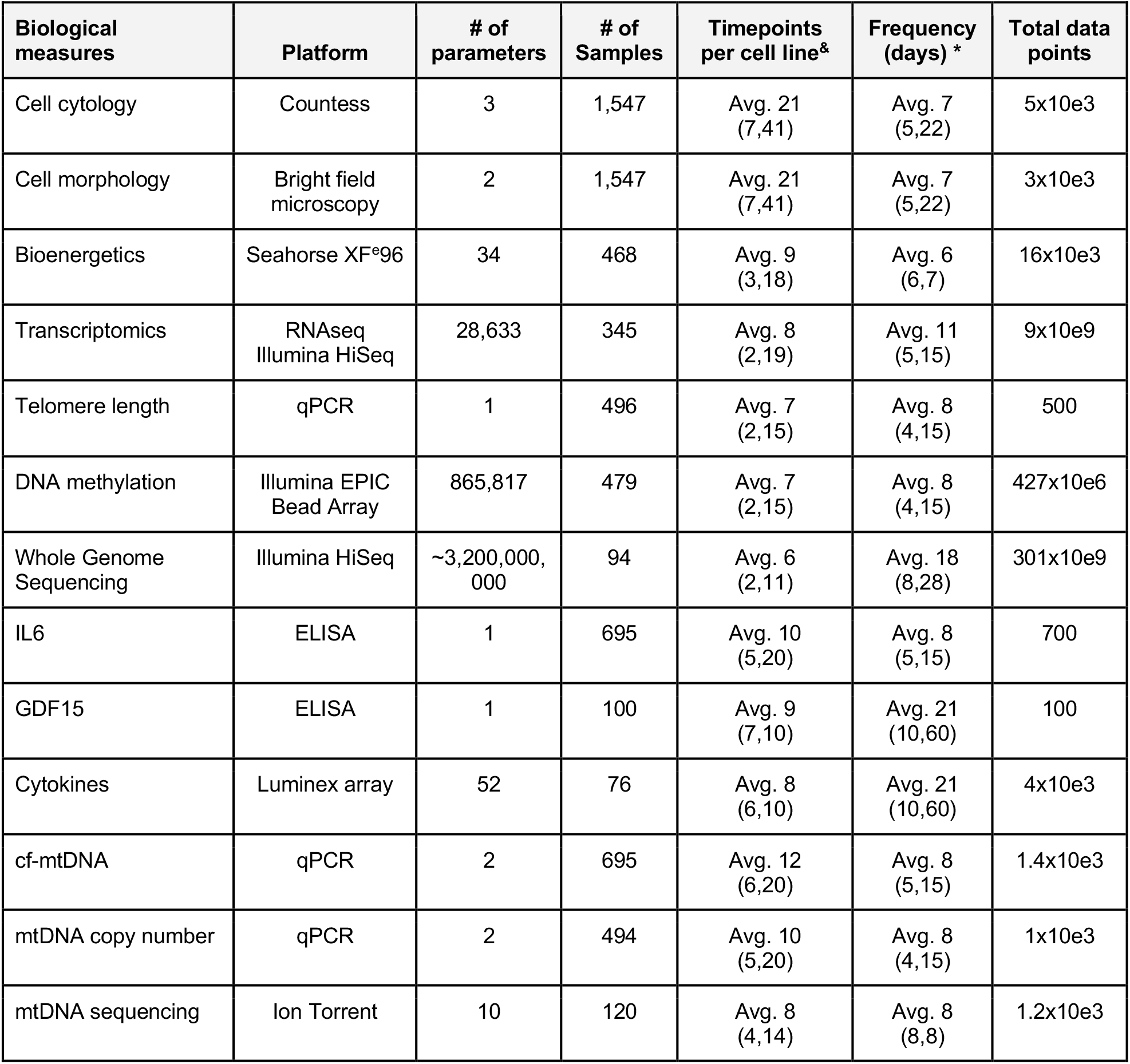
Dimensionality of biological measures available in this dataset. Molecular, cellular and bioenergetic features, the technique or platform used to generate the data, their temporal resolution, and the dimensionality of datasets for each biological measure. ^&^ Values are averages with minimum and maximum number of timepoints (min, max). * The frequency of passaging differs by study Phase: Phase I, 5 days; Phase II, 5 days; Phase III, 7 days; Phase IV, 7 days; Phase V, 5 days.

**Table 4** lists the five main phases for which the study was conducted (Phases I-V). The time periods and experimenters associated with each study phase.

**Table 4.** Study phases, cell lines used, and treatments applied in the *Cellular Lifespan Study*. Five phases conducted by different investigators with varying study lengths, cell lines, treatments, and exposures.

| Study Phase | Investigator | Time period | Cell lines | Treatments |
| --- | --- | --- | --- | --- |
| Phase I | G.S. | Oct 2017-Feb 2018 (4 mo) | HC5,6 | 1,2,6,11,12,13,19,20,21,22 |
| Phase II | G.S. | Aug 2018-May 2019 (10 mo) | HC1,2,3,<br>SURF1_1,2,3 | 1,2,3,4,10,14,15,18 |
| Phase III | G.S. | Feb 2019-April 2019 (3 mo) | HC1,2,4 | 1,2,5,6,16,17 |
| Phase IV | J.M. | May 2019-Oct 2019 (6mo) | HC1,2 | 1,7,8,9 |
| Phase V | A.S.M. | Oct 2020-June 2021 (8mo) | HC1,2,3,<br>SURF1_1,2,3 | 1,5 |

### Experimental treatments

1. **Control (untreated)**
  a. **Rationale:** Cells are grown under metabolically diverse conditions of Dulbecco’s modified Eagle’s medium (DMEM, Thermofisher #10569044) containing 5.5 mM glucose, 1 mM pyruvate, 4 mM GlutaMAX™, supplemented with 10% fetal bovine serum, 1% non-essential amino acids, 50 mg/ml Uridine, and 10 mM Palmitate.
  b. **Design & dose:** No treatment. To enable direct comparison of control growth trajectories to treatments (diluted in the vehicle DMSO), untreated controls were grown in 0.001% DMSO (Sigma-Aldrich #D4540).
  c. **Duration:** 0-270 days
  d. **Study Phase**: I-V
2. **Chronic Dexamethasone (DEX)**
  a. **Rationale:** Glucocorticoid receptor agonist, activates transcription of >1,000 genes ^58^. Used as a mimetic of chronic neuroendocrine or psychosocial stress in animal studies ^59^. This treatment was used to examine the effects of chronic activation glucocorticoid signaling, a major evolutionary conserved stress pathway, on aging- and metabolism-related processes.
  b. **Design & dose:** (PubChem CID: 5743, SID: 46508930) Chronic 100nM (in EtOh) dose every passage, Sigma-Aldrich #D4902.
  c. **Duration:** 20-270 days (chronic)
  d. **Study Phase**: I-III
3. **Oligomycin (Oligo)**
  a. **Rationale:** Inhibition of the mitochondrial OXPHOS system by inhibiting the ATP synthase (Complex V). Oligo treatment causes depletion of mitochondria-derived ATP, hyperpolarization of the membrane potential, and triggers retrograde signaling that activates the integrated stress response (ISR) ^60^. This treatment was used to inhibit OXPHOS downstream from the respiratory chain, and thereby examine the effect of chronic OXPHOS dysfunction.
  b. **Design & dose:** Chronic, 1nM (stored in DMSO) dose every passage, Sigma-Aldrich #75351
  c. **Duration:** 20-220 days
  d. **Study Phase**: II
4. **Mitochondrial Nutrient Uptake Inhibitors (mitoNUITs)**
  a. **Rationale:** Inhibiting the import of three major substrates into mitochondria, including i) *pyruvate*, with UK5099, an inhibitor of the mitochondrial pyruvate carrier (MPC); ii) *fatty acids*, with Etomoxir, an irreversible inhibitor of carnitine palmitoyltransferase-1 (CPT-1) that prevents the transport of fatty acyl chains from the cytoplasm to the mitochondria; and iii) *glutamine*, with BPTES, an inhibitor of glutaminase GLS1 that converts glutamine to glutamate inside the mitochondria. This treatment was used to starve the tricarboxylic acid (TCA) cycle of carbon intermediates, thus inhibiting OXPHOS upstream of the respiratory chain.
  b. **Design & dose:** UK5099 (PubChem CID: 6438504, SID: 329825569) chronic at 2μM (in DMSO), Sigma-Aldrich #PZ0160; Etomoxir (PubChem CID: 123823) chronic at 4μM (in ddH_2_O), Sigma-Aldrich #E1905; BPTES (PubChem CID: 3372016) chronic at 3μM (in DMSO), Sigma-Aldrich #SML0601; dosed every passage.
  c. **Duration:** 20-210 days
  d. **Study Phase**: II
5. **Hypoxia**
  a. **Rationale:** Oxygen is the terminal electron acceptor at respiratory chain complex IV (cytochrome c oxidase), and decreasing ambient oxygen tension from 21% to 3% has been shown to influence cellular bioenergetics. In cellular and animal models of mitochondrial disease, including the Ndufs4 deficient mouse ^30^, hypoxia treatment has shown promise to alleviate the disease phenotype and extend lifespan. In relation to cellular aging, 3% oxygen tension also extends cellular lifespan in murine and human fibroblasts ^61–65^.
  b. **Design & dose:** Cells chronically grown at 3% O_2_ (5% CO_2_), except periodic exposure to ambient 21% O_2_ during passaging (∼3 hours, once each week). This experiment was run twice (Study Phase III and V). Phase III compared healthy to chronic DEX cells in hypoxia, while Phase V compared healthy to SURF1-mutant cells in hypoxia. Additionally, all seahorse bioenergetic measurements were measured in 21% oxygen (Phase V included an overnight incubation at 21% O_2_).
  c. **Duration:** 0-70 days
  d. **Study Phase**: III & V
6. **Contact Inhibition**
  a. **Rationale:** Allowing cells to fill up the dish and enter a quiescent state allows for the experimentally untangle the role of cell division in time-dependent changes. Contact-inhibited fibroblasts continue to undergo morphological changes, exhibit >88% reduced division rate on average, and display skin-like tissue appearance after months in culture.
  b. **Design & dose:** After thawing and adjustment to culture environment cells are plated in multiple flasks at high density with marked collection points with the 0 time point collected 7 days after the initial plating. Media is changed weekly, at the same time points as dividing cells are passaged.
  c. **Duration:** 20-140 days
  d. **Study Phase**: III
7. **Galactose**
  a. **Rationale:** Galactose is a non-fermentable sugar and its oxidation into pyruvate through glycolysis yields no ATP, thereby forcing cells to rely solely on OXPHOS for ATP production ^66^.
  b. **Design & dose:** Galactose (PubChem CID: 6036, SID: 178101363) chronic at 5.5mM (stored in ddH_2_O) dose every passage, Sigma-Aldrich #G5388.
  c. **Duration:** 20-170 days
  d. **Study Phase**: IV
8. **2-deoxy-d-glucose (2-DG)**
  a. **Rationale:** 2-deoxy-d-glucose (2DG) is an inhibitor of hexokinase ^67^ that blocks metabolic flux and ATP synthesis from the metabolism of glucose through glycolysis.
  b. **Design & dose:** 2-DG (PubChem CID: 108223, SID:24893732) chronic at 1mM (in ddH_2_O) dose every passage, Sigma-Aldrich #D3179.
  c. **Duration:** 20-170 days
  d. **Study Phase**: IV
9. **Beta-hydroxybutyrate (and 0mM glucose)**
  a. **Rationale:** Caloric restriction has been shown to extend lifespan. Beta-hydroxbutyrate is a ketone body that is induced in caloric restriction and acts as a signaling metabolite to effect gene expression in diverse tissues ^68^.
  b. **Design & dose:** hydroxybutyrate (PubChem CID: 10197691, SID: 57651496) chronic at 10mM (in ddH_2_O) dose every passage, Sigma-Aldrich #54965.
  c. **Duration:** 20-170 days
  d. **Study Phase**: IV
10. **5-azacytidine (5-aza)**
  a. **Rationale:** 5-aza was used to induce global demethylation of the genome, thus testing if direct alteration of the methylome would reset the aging of our cells. Note, no significant change was seen in the growth rate or global DNA methylation after treatment, suggesting that the dose used may have been insufficient to induce robust alterations in DNA methylation.
  b. **Design & dose:** 5-aza (PubChem CID: 9444, SID: 24278211) acute treatment for 2 passages (∼10days) at 1μg/mL (stored in PBS), Sigma-Aldrich #A2385.
  c. **Duration:** 60-230 days
  d. **Study Phase**: II
11. **MitoQ**
  a. **Rationale:** MitoQ is a mitochondria-targeted antioxidant compound used specifically in the mitochondrial compartment ^69^. MitoQ was used to test how reducing mitochondrial oxidation would influence cellular aging, or moderate the influence of chronic stressors (see *Treatment 19*).
  b. **Design & dose:** MitoQ (PubChem CID: 11388332, SID:134224101) chronic treatment at 10nM (stored in DMSO), provided by author M.P.M.
  c. **Duration:** 20-120 days
  d. **Study Phase**: I
12. **N-Acetyl-L-cysteine (NAC)**
  a. **Rationale:** NAC acts as a precursor of glutathione (GSH), which is itself a direct antioxidant and a substrate for antioxidant enzymes ^70^, NAC also generates H2S and sulfanes that can act as antioxidants ^71^. NAC was used to test whether reducing total cellular oxidative stress burden by scavenging of reactive oxygen species may influence cellular aging, or moderate the effect of chronic stress (see *Treatment 20*).
  b. **Design & dose:** NAC (PubChem CID: 12035, SID: 24277970) chronic treatment at 2mM (in ddH_2_O), Sigma-Aldrich #A7250.
  c. **Duration:** 20-120 days
  d. **Study Phase**: I
13. α-ketoglutarate (α-KG)
  a. **Rationale:** α-KG (also 2-oxoglutarate) is a key tricarboxylic acid cycle (TCA) cycle metabolite that is a substrate for 2-oxoglutarate-dependent dioxygenases (2-OGDD), and necessary cofactor for enzymes that perform demethylation of proteins and DNA ^72^. α-KG was used to shift the α-KG-to-succinate ratio, which was hypothesized to promote DNA demethylation ^73^. The cellular uptake and bioavailability of α-KG was not monitored.
  b. **Design & dose:** A-ketoglutarate (PubChem CID: 164533, SID: 329766750) chronic treatment at 1mM (in ddH_2_O), Sigma-Aldrich #75890.
  c. **Duration:** 20-130 days
  d. **Study Phase**: I
14. **Oligomycin + DEX**
  a. **Rationale:** The complex V inhibitor oligomycin was used in combination with DEX to examine their interactions. DEX increases mitochondrial OXPHOS-derived ATP production, which is inhibited by Oligo, suggesting that DEX and Oligo may have antagonistic effects.
  b. **Design & dose:** same as Treatments 2 and 3.
  c. **Duration:** 20-70 days
  d. **Study Phase**: II
15. **mitoNUITs + DEX**
  a. **Rationale:** Because the increase in mitochondria ATP production induced by DEX requires the uptake of carbon substrates, which is inhibited by mitoNUITs. Both treatments were used in parallel to examine if mitoNUITs would interfere with the effects of DEX on cellular bioenergetics and signaling.
  b. **Design & dose:** same as Treatments 2 and 4.
  c. **Duration:** 20-270 days
  d. **Study Phase**: II
16. **Hypoxia + DEX**
  a. **Rationale:** Hypoxia triggers a shift towards glycolytic metabolism, whereas DEX causes a shift towards OXPHOS. Both treatments were used in parallel to examine if DEX hypoxia could rescue the chronic effects of DEX on energetic parameters and aging markers.
  b. **Design & dose:** same as Treatments 2 and 5.
  c. **Duration:** 20-70 days
  d. **Study Phase**: III
17. **Contact inhibition + hypoxia**
  a. **Rationale:** Both contact inhibition and hypoxia reduce cellular division rate, and partially recreate some of the natural conditions of skin fibroblasts in the human body. Both were used in parallel as an attempt to recapitulate as closely as possible *in vivo* conditions and evaluate the influence of this state on aging markers.
  b. **Design & dose:** same as Treatments 5 and 6.
  c. **Duration:** 20-140 days
  d. **Study Phase**: III
18. **5-azacytidine + mitoNUITs**
  a. **Rationale:** Here we tested the idea that mitochondrial activity stores the memory of epigenetic state. By demethylating the genome with 5-azacytidine and then simultaneously diverting energy away from the ETC we hypothesized that the genome would take longer to be remethylated back to its original state.
  b. **Design & dose:** same as Treatments 4 and 10.
  c. **Duration:** 60-230 days
  d. **Study Phase**: II
19. **MitoQ + DEX**
  a. **Rationale:** DEX causes a bioenergetic shift towards OXPHOS, and causes premature aging based on several biomarkers (unpublished). MitoQ was used to examine if these effects of chronic glucocorticoid stimulation could be alleviated by buffering mitochondrial ROS, which would suggest that part of the accelerated aging phenotype in DEX-treated cells is in part driven by mitochondrial ROS.
  b. **Design & dose:** same as Treatments 2 and 11.
  c. **Duration:** 20-130 days
  d. **Study Phase**: I
20. **NAC + DEX**
  a. **Rationale:** Similar to *Treatment 19*, NAC was used in parallel with DEX to examine if the chronic effects of glucocorticoid signaling could be alleviated by buffering of ROS in the cytoplasmic compartment (vs MitoQ, for mitochondrial ROS).
  b. **Design & dose:** same as Treatments 2 and 12.
  c. **Duration:** 20-120 days
  d. **Study Phase**: I
21. **Pulsated DEX**
  a. **Rationale:** This treatment was used to examine the effects of a more physiological pulsatile activation of glucocorticoid signaling, compared to its chronic activation (Treatment 2).
  b. **Design & dose:** Same molecular composition as chronic DEX (100nM, see Treatment 2). Cells were treated once, for 30 min, before each passage (every 5-7 days).
  c. **Duration:** 20-120 days (pulsated)
  d. **Study Phase**: I
22. **Contact Inhibition & regrowth**
  a. **Rationale:** Allowing cells to continue their replicative lifespan after being held in contact inhibition for several weeks allows for the experimental determination of whether cultured cells remember their divisional age and contain the same total replicative lifespan.
  b. **Design & dose:** After 80 days of contact inhibition cells were allowed to continue dividing until replicative exhaustion (i.e. ‘Contact Inhibition Regrowth’).
  c. **Duration:** 80-210 days
  d. **Study Phase**: I

### Technical Validation

The number of days per passage were systematically recorded (and can be derived from the ‘*Date_time_of_passage*’ variable, for each cell count timepoint in the database). Variation and deviations in the number of days between passages increase towards the latter part of the lifespan for all lines and treatments because of the limited number of cells once enter quiescence and senescence (**Figure 5A**).

**Figure 5.**
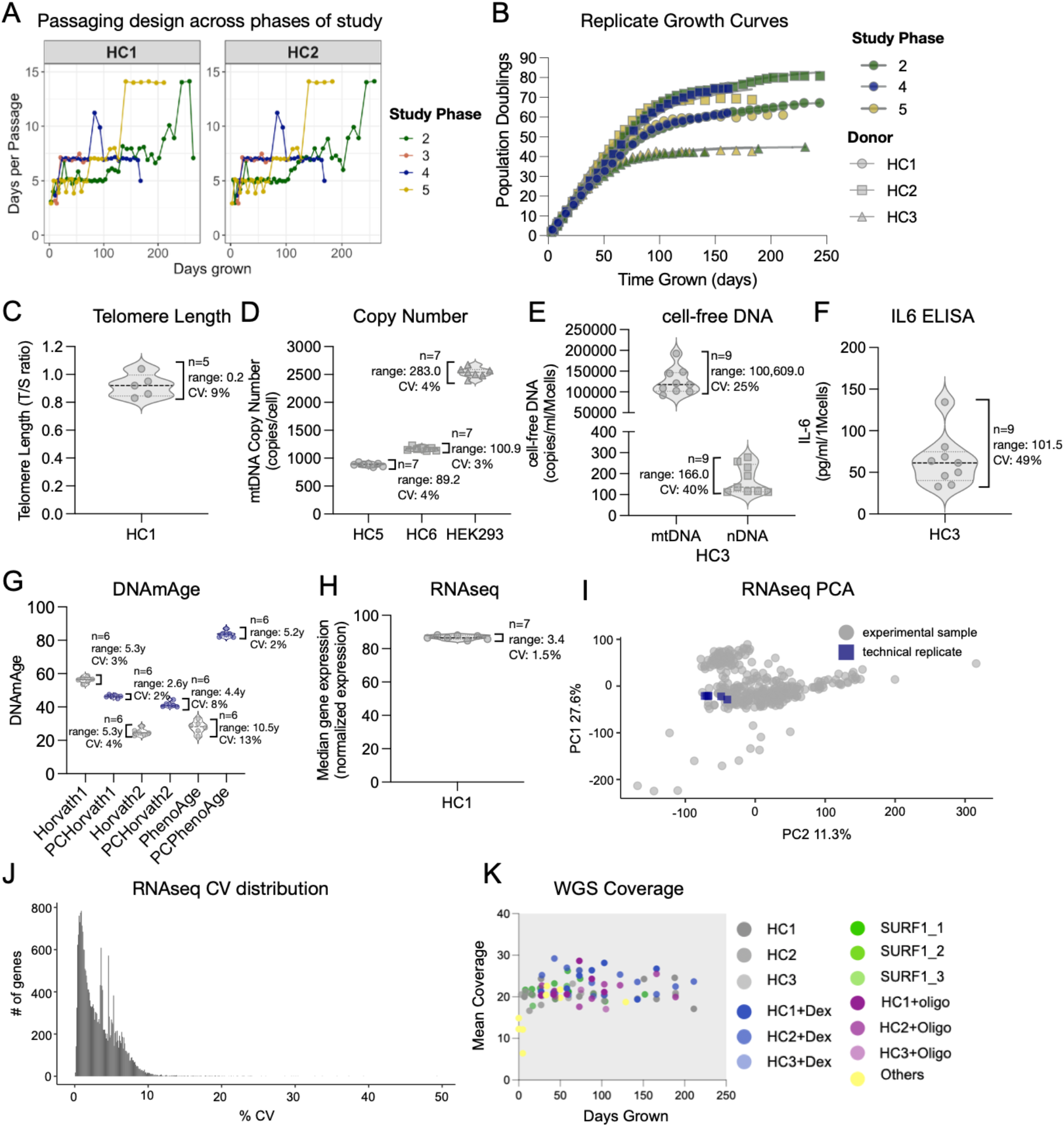
Experimental replication and technical variation across multi-modal measurements. (**A**) Timecourse of the number of days between each passage for each phase of the study. (**B**) Growth curve of donors repeated at different phases of the study. Color indicates phase of study and shape indicate cell line donor. (**C-H**) Variation in technical replicates for (**C**) telomere length, (**D**) mtDNA copy number, (**E**) cell-free DNA, (**F**) IL6 as measured by ELISA, (**G**) DNAmAge as estimated by three epigenetic clock and there PC-adjusted calibrations, and (**H**) average RNAseq-based transcript levels across all expressed genes. (**I**) Principal component analysis of all expressed genes for the full 354 sample RNAseq dataset. Technical replicates are highlighted in blue (n=7). (**J**) Coefficient of variations in transcript per million (TPM) across the 7 RNAseq technical replicates for each mapped gene. The majority of gene have CVs < 5%, and 99% of genes have CVs < 10%. (**K**) Mean sequencing coverage for whole genome sequencing (n=94 samples).

Two of the 6 healthy controls’ growth curves were repeated 4 times, over a ∼2 year period, in separate phases by different experimenters (see **Table 4**), confirming that the growth curves were reproducible (**Figure 5B**). Chronic DEX (Treatment 2) was repeated in study Phases I, II, and III. Hypoxia (Treatment 5) was repeated in study Phases III and V. SURF1-mutant cells were run in both study Phases II and V. All data reflecting independent growth curves on multiple parameters can be visualized on our webtool (https://columbia-picard.shinyapps.io/shinyapp-Lifespan_Study/) and from the downloaded data.

The telomere length qPCR assay contained 5 technical replicates of the same sample (HC1, passage 10, 42 days grown). Replicates had a C.V. of 9% (range of 0.21 T/S units) (**Figure 5C**).

The mtDNA copy number qPCR assay contained 7 technical replicates across 7 plates of the same 2 samples (HC5 & HC6). Replicates had an average C.V. of 3.5% (range: 95.05 copies/cell) (**Figure 5D**).

The cell-free DNA qPCR assay contained 9 technical replicates across 8 plates of the same samples (HC3, passage 32, 189 days grown). An aged sample was selected to ensure robust extracellular DNA levels. Replicates of cf-mtDNA and cf-nDNA measurements had an average C.V. of 25% and 40% (range: 100,609 and 166 copies/mL/10^6^ cells, respectively) (**Figure 5E**).

The IL6 ELISA assay contained 9 technical replicates across 8 plates of the same samples (HC3, passage 32, 189 days grown). An old media sample (robust extracellular IL6 levels) was selected as a positive control. Replicates had a C.V. of 49% (range: 101.5 pg/mL/10^6^ cells) (**Figure 5F**).

The technical variation in DNAm on the EPIC array was also previously determined on six DNA replicates from HC5 (GEO #GSE131280). DNAmAge computed from the combination of multiple probes or CpG sites showed a coefficient of variation, for selected clocks, of 3% for the Horvath1 (PanTissue) clock, 4% for the Horvath2 (Skin&Blood) clock, and 13% for the PhenoAge clock (**Figure 5G**). After PC-adjustment based on ^51^, the technical variation between samples was reduced to 2%, 8%, and 2%, respectively, indicating moderately robust technical validity at the single-CpG level, but high validity when multiple CpGs are combined into multivariate DNAm clocks. The DNAm dataset also contains 3 replicate longitudinal experiments of HC1 & HC2 that can be used to quantify the variability in the longitudinal rates of epigenetic aging using the investigator’s preferred clock(s) or individual CpGs.

Technical variation for RNA- and DNA-based OMICS measures with >100 samples (i.e. RNAseq, DNAm, telomere length etc.) was determined by distributing the same biological sample (healthy HC1 fibroblasts, untreated, passage 6, 21 days grown) across multiple plates and sequencing lanes. For RNAseq, the technical assay variation was determined from 7 biological replicates. The average coefficient of variation (C.V., standard deviation divided by the mean) in normalized read counts across all genes was 1.47% (range = 3.42 normalized expression units (**Figure 5H**). **Figure 5J** shows the frequency distribution of C.V. for each mapped gene, and how technical variation for individual genes influenced sample position on a 2-component principal component analysis (PCA) across all study samples, indicating good reliability (**Figure 5I**).

The mean sequencing depth for WGS was >20X for most samples, as shown in **Figure 5k**.

### Usage Notes

Here we mention some of the limitations of the dataset that should be considered for the analysis and interpretation of findings.

#### Sample Exclusion

All samples and measurements that did not pass quality control, regardless of the assay, were excluded from the published dataset (i.e. ‘Supplemental File 1’ & ShinyApp). See ‘Data Access’, section for access to raw data files.

#### Prestudy passaging

Fibroblast cell lines were sourced from different vendors and isolated using varying culture methods (**Table 1**). Additionally, donor medical history, chronological age, and environmental exposures add additional variability between cell lines. Due to these constraints, it is not possible to determine the exact number of population doublings cells underwent before performing the study. These constraints could influence the cumulative population doublings of each cell line but should not affect rates of aging or age-related trajectories.

#### Pre-study freeze-thaw cycles

Study Phases I and II involved a single freeze-thaw cycle in liquid nitrogen in our laboratory from the cell obtained; while an additional cycle occurred in cells used in Phases III, IV, V.

#### Normalization to cell numbers

The raw values for some features are influenced by the number of cells in the culture flask at each passage. For example, secretome data that includes extracellular levels of cytokines or cell-free mitochondrial DNA (cf-mtDNA) in the cell media are determined not only by the secretion rate, but also by the total number of cells contributing to the signal in the flask. Therefore, secreted factors levels are normalized to cell number at the time of harvesting media, which represents the amount of analyte released per cell counted. To obtain raw concentrations media concentrations, the investigator can multiply the normalized values (*variable_name_example* e.g. ‘*IL6_ELISA_Upg_per_ml’*) by the cell count (*variable_name_in_database_for_cellcount, e*.*g. ‘Cells_counted_UmillionCells’*) at each time point. Cell counts should include the fraction of dead cells at any given passage unless otherwise established secretion from exclusively live cells.

#### Cell seeding density

In Phase V of the study, we improved our calculation of seeding numbers to ensure <80% confluency at each passage. This minor change in protocol could contribute to differences between replicate experiments and trajectories of metabolic rate and other potential differences between Phase V and earlier phases.

#### Sample size and robustness of experimental treatments

As detailed in Table 2, this *Cellular Lifespan* dataset includes longitudinal assessments of treatment conditions in primary human fibroblasts from multiple independent donors, in some cases replicated multiple times. This is the case for untreated (Treatment #1), chronic DEX (#2), and hypoxia (#5) treatments. These time-series data enable robust modeling of time-related dynamics and treatment effects, using the user’s preferred methods. Other treatment conditions were either performed in a smaller number of donors, particularly HC1 (male) and HC2 (female), and/or for some treatments the timecourse experiment was not subsequently repeated. These conditions should be considered exploratory and may serve as preliminary data for subsequent experiments. All treatment conditions were conducted in at least two different donors, and each experiment contained multiple timecouse data points. The more limited domains of the dataset enable to calculate robust estimates of effect sizes, directionality of effects, and effects across two donors of different biological sex.

#### Rate of epigenetic aging

To obtain stable estimates of epigenetic aging without potential artifacts attributable to the early effects of culture or before the onset of treatments, or to later alterations in the epigenome (i.e., non-linear trajectories on DNAmAge clocks) related to quiescence or senescence towards the end of life, rates of epigenetic aging were determined by taking the linear slope for each cell line from 25 to 75 days of growth.

#### Future analyses

This dataset contains multi-modal data that can be used to investigate each of the hallmarks of aging (see **Figure 4**). In particular, the high-resolution timecourses are ideal to model age-related trajectories with either linear or complex non-linear functions. These models could then be systematically classified and statistically examined using functional regression approaches, for example, to identify (groups of) parameters exhibiting similar related age-associated trajectories. Such parameters would indicate co-regulation and could inform subsequent mechanistic studies aiming to establish causal pathways, bioenergetic parameters, enzymes, or genes that drive specific aging trajectories. Because the average timecourse contains ∼12 timepoints, and because interpolation likely overfits beyond the 3-sample resolution rule-of-thumb, non-linear modeling efforts should be limited to 4 inflection points across the cellular lifespan.

It is also possible to leverage the repeated measures design across portions of the lifespan, or across the whole lifespan, to examine stable (i.e. time-invariant) differences between treatment groups. Examples include: i) the SURF1 mutant cells effects relative to control, which triggers hypermetabolism, a robust hypersecretory phenotype, the transcriptional integrated stress response, and accelerates several markers of cellular aging ^74^, and ii) the chronic Dex treatment effects on control fibroblasts, which alters cytological, transcriptional, secretory, and (epi)genomic aging markers ^75^.

Finally, these longitudinal data can be used to develop or validate new penalized regression algorithms or “epigenetic clocks” ^51^.

## Data Access

All data can be accessed, visualized, and downloaded without restrictions at https://columbia-picard.shinyapps.io/shinyapp-Lifespan_Study/. This simple interface allows users to select (and de-select) the donors of interest, select the experimental treatment(s), and visualize the time course data in an realtime-updatable display panel (**Figure 6**). Users can select multiple donors and-or treatments simultaneously to visualize the effects of interest and explore the data. All data visualized is downloadable as a .csv file which can further be found directly at: https://figshare.com/articles/dataset/Lifespan_Study_Data/18441998. The app will be regularly updated with new data as additional lifespan experiments and analyses are performed.

The unprocessed RNAseq (GSE179848) and EPIC DNA methylation array data (GSE179847) can also be accessed and downloaded in full through Gene Expression Omnibus (GEO). WGS coverage and variant information can be accessed on the ShinyApp, and the complete data is available upon request. Brightfield microscopy images can be downloaded at: https://figshare.com/articles/dataset/Brightfield_Images_for_Cellular_Lifespan_Study/18444731. Raw Seahorse assay files along with corresponding data analysis scripts can be found at : https://github.com/gav-sturm/Cellular_Lifespan_Study/tree/main/Seahorse.

**Figure 6.**
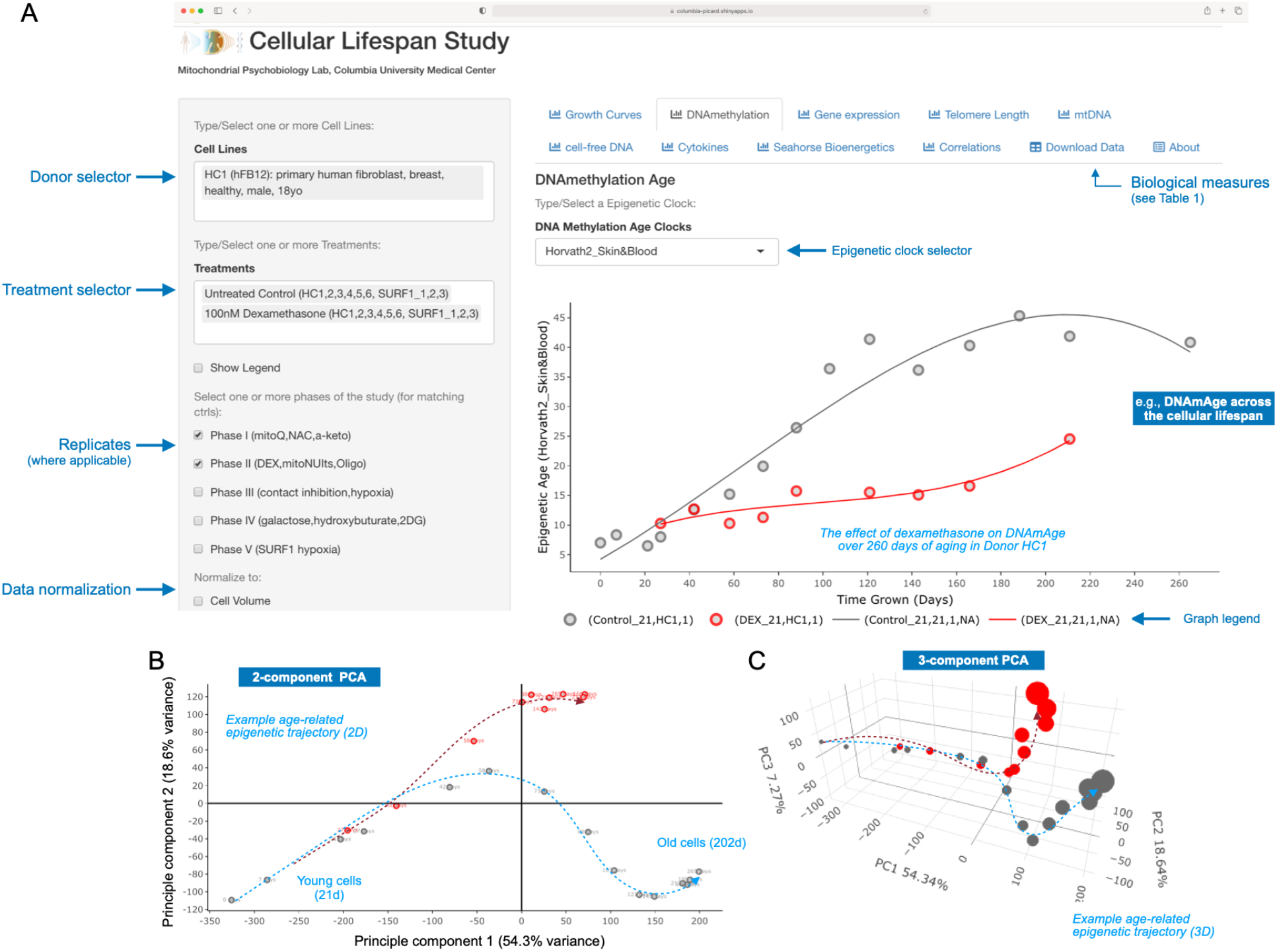
Data visualization and exploration on the Cellular Lifespan ShinyApp. (**A-C**) Visualization example of high-dimensionality epigenetic DNA methylation (DNAm) data using tools on the ShinyApp. (A) Interactive timecourse of the DNAm Skin&Blood clock (Horvath2, a multivariate algorithm trained using penalized elastic net regression) across the cellular lifespan of HC1 fibroblasts, both untreated (grey) and treated chronically (red) with 100nM of dexamethasone (DEX, glucocorticoid receptor agonist) to mimic chronic stress exposure. Note that the x axis represents time in culture, which can be changed using the selector menu on the left to “population doublings” to take into account the reduced division rate in Dex-treated cells. Other biological measures are visualized via tabs on the top. (**B-C**) Interactive principal component analysis of the 45,000 significant age-related CpGs in 2D (B) and 3D (C). The frequency and duration of different cytological, molecular, and bioenergetic measurements vary by cell lines and experimental conditions. See Figure 3, Tables 2-4, and Supplemental File 2 for details. The Shiny App can be accessed, and the data downloaded, at https://columbia-picard.shinyapps.io/shinyapp-Lifespan_Study.

## Code Availability

Code is available at https://github.com/gav-sturm/Cellular_Lifespan_Study

## Acknowledgments

This work was supported by NIH grant AG066828.

## Author contributions

G.S. and M.P. designed the study. M.H. contributed cell lines. G.S., A.S.M., J.M. performed cell culture and sample collection. K.R.K contributed cytokine array and whole genome sequencing data. J.L. contributed telomere length data. V.P. and C.B. contributed mtDNA sequencing data. B.A.K. and S.A.W. contributed cell-free DNA data. S.H., M.L., A.H.C., D.B., S.W. contributed DNAm algorithms. G.S. curated and organized the dataset with A.S.M. G.S. developed the ShinyApp. G.S. and M.P. drafted the manuscript, with revisions from D.B. All authors reviewed and approved the final version of this manuscript.

## Competing interests

The authors declare no competing interest.

## Supplementary Files

Supplementary File 1. Cellular Lifespan Study complete dataset.

Supplementary File 2. Heatmaps of available experimental data.

## Notes

https://columbia-picard.shinyapps.io/shinyapp-Lifespan_Study/

https://www.ncbi.nlm.nih.gov/geo/query/acc.cgi?acc=GSE179849

https://figshare.com/articles/dataset/Lifespan_Study_Data/18441998

https://figshare.com/articles/dataset/Brightfield_Images_for_Cellular_Lifespan_Study/18444731

https://github.com/gav-sturm/Cellular_Lifespan_Study/

### Competing Interest Statement

The authors have declared no competing interest.

### Summary of Updates

Fixed the resolution on the figures for improved visibility.

