## Supplemental File 2 for "A Multi-Omics Longitudinal Aging Dataset in Primary Human Fibroblasts with Mitochondrial Perturbations"

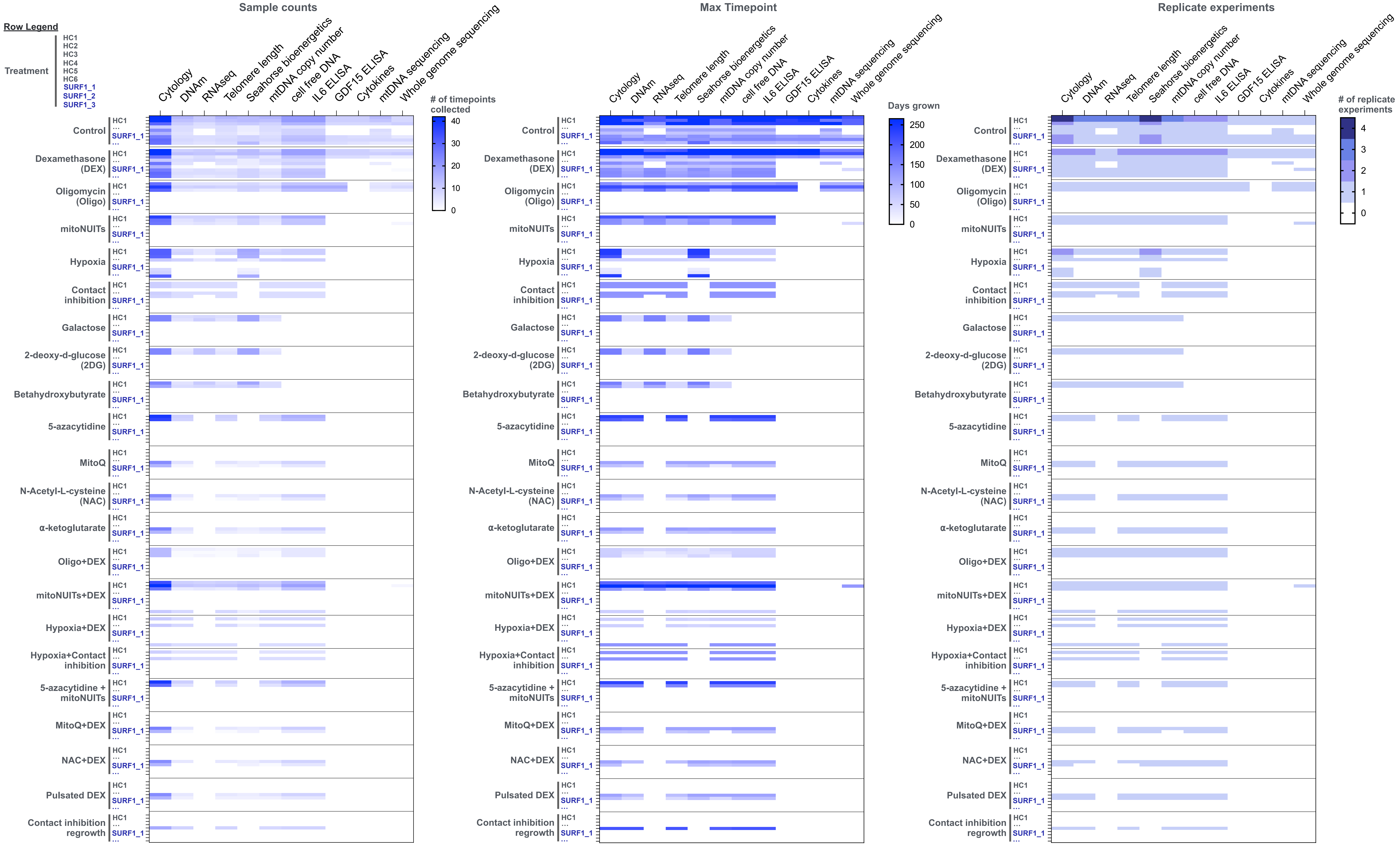


**Supplemental File 2. Sample counts, max timepoint, and replicated experiment counts for each measured parameter.** (**A**) Heatmap of the number of timepoints for each parameter (columns) associated with a given donor cell line and treatment (rows). (**B**) Heatmap of the maximum timepoint collected for each parameter (columns) associated with a given donor cell line and treatment (rows). The highest value is 265 days. The maximal duration of each experiment was determined by either the absence of divisions for a 30-day period, reaching the target endpoint for a specific experiment (e.g., Treatment 5: Hypoxia), or rare events of bacterial contamination. (**C**) Heatmap of the number of repeat experiments (i.e. Study Parts) for each parameter (columns) associated with a given donor cell line and treatment (rows).
